# mTOR regulates Wnt signaling to promote tension-mediated lens vesicle closure

**DOI:** 10.1101/2025.02.24.639869

**Authors:** Qian Wang, Hao Wu, Yingyu Mao, Alyssa Chow, Michael Bouaziz, Yihua Wu, Xin Zhang

**Affiliations:** Department of Ophthalmology and Visual Sciences, Washington University School of Medicine, St. Louis, MO 63110, USA; Departments of Ophthalmology, Pathology and Cell Biology, Columbia University, New York, NY 10032, USA

**Keywords:** Peters anomaly, ciliary margin, lens stalk, mTOR, Wnt3, Rac1

## Abstract

Lens vesicle closure is a pivotal event in ocular morphogenesis, and its disruption underlies Peters anomaly, a leading congenital cause of corneal opacity. Here, we elucidate a mechanistic hierarchy in which mTOR-Wnt signaling orchestrates cytoskeletal tension to drive this process. Conditional ablation of mTOR in the lens ectoderm induces aberrant corneal-lenticular stalk formation and transdifferentiation of the ciliary margin into neural retina. mTOR inhibition suppresses Wnt3 expression, and Wnt3 displayed a similar lens stalk phenotype, positioning mTOR as an upstream regulator of Wnt ligand production. Complete ablation of lens-derived Wnt ligands via deletion of the Wnt transporter Wls exacerbates developmental defects, triggering anterior lens herniation and ciliary margin development failure. Disruption of β-catenin-mediated Wnt signaling or dual deletion of Wnt co-receptors Lrp5/6 in lens ectoderm similarly prevents vesicle closure, recapitulating lens herniation. Strikingly, Rac1 deletion rescues corneal-lenticular stalk phenotypes in mTOR, Wls, and β-catenin mutants, directly linking Wnt effectors to cytoskeletal remodeling. Our findings establish an mTOR-Wnt-Rac1 signaling axis as the core regulator of cytoskeletal tension required for lens vesicle closure.

## Introduction

Mammalian lens development is a tightly orchestrated process involving precise spatiotemporal coordination of cell fate specification, morphogenetic movements, and tissue separation (Cvekl and Zhang, 2017). In mice, this cascade initiates at embryonic day 9.5 (E9.5) as the lens ectoderm (LE) thickens to form the lens placode. By E10.5, the placode undergoes invagination, creating a transient lens vesicle that must subsequently detach from the surface ectoderm—a critical step enabling divergent differentiation: the vesicle forms the transparent crystalline lens, while the adjacent ectoderm gives rise to the cornea. Both tissues share an absolute requirement for optical clarity, yet their developmental paths diverge irreversibly. Failure of lens-corneal separation disrupts this trajectory, resulting in Peters anomaly type II (PA2), a frequent cause of congenital corneal opacity (Bhandari et al., 2011; Nischal, 2012). PA2 is notoriously refractory to treatment, with corneal transplantation often yielding suboptimal visual outcomes. Thus, understanding the etiology of Peters anomaly is critical not only for revealing the cellular biology underlying lens-cornea separation but also for understanding the origins of corneal opacity and the generally poor prognosis in these patients.

Canonical Wnt signaling is initiated by Wnt ligand binding to Frizzled (Fzd) and Lrp5/6 co-receptors, triggering β-catenin stabilization and nuclear translocation to activate target gene transcription (Clevers and Nusse, 2012). Intriguingly, LE-specific β-catenin deletion generates ectopic mini-lenses outside the lens territory, while constitutive β-catenin activation suppresses endogenous lens induction (Smith et al., 2005). These findings suggest that precise spatial regulation of Wnt activity is critical for lens fate specification, yet β-catenin’s dual roles in cell adhesion at adherens junctions and Wnt-mediated transcriptional regulation complicate mechanistic interpretation. Temporal studies reveal that β-catenin remains indispensable post-induction: its elimination after lens vesicle formation blocks lens fiber differentiation (Cain et al., 2008), indicating stage-specific requirements for Wnt signaling. Conversely, systemic *Lrp6* knockout or LE-specific ablation of the Wnt chaperone *Wls* causes anterior lens herniation (Carpenter et al., 2015; Stump et al., 2003). These phenotypes point to an autocrine Wnt signaling role in maintaining lens positional integrity, though the downstream effectors mediating this process remain undefined. Notably, *Wls* deletion also impairs ciliary margin (CM) and retinal pigment epithelium (RPE) development, implicating LE-derived Wnt ligands in broader optic cup patterning (Balasubramanian et al., 2021). Together, these studies underscore Wnt signaling as a multifaceted regulator of ocular morphogenesis, with context-dependent roles in cell fate determination, structural maintenance, and tissue-tissue communication.

The mechanistic target of rapamycin (mTOR) kinase operates as the catalytic core of two functionally distinct complexes, mTORC1 and mTORC2, defined by their unique regulatory subunits Raptor and Rictor, respectively (Laplante and Sabatini, 2012). These complexes act as central signaling hubs, integrating nutrient availability, energy status, and growth signals to coordinate cellular metabolism with developmental programs. mTORC1 primarily governs anabolic processes, including protein synthesis via phosphorylation of downstream effectors 4EBP1 and S6K, while simultaneously repressing catabolic pathways such as autophagy. In contrast, mTORC2 phosphorylates AKT at serine 473, a post-translational modification essential for full AKT activation—a key regulator of survival and growth signaling. Evolutionarily conserved across eukaryotes, the mTOR pathway exemplifies a fundamental mechanism linking metabolic homeostasis to cellular growth. During development, this nexus is particularly critical, as morphogenetic processes demand precise spatiotemporal control of energy-intensive events like cytoskeletal remodeling and tissue patterning. By synchronizing metabolic states with signaling cascades, mTOR complexes ensure that cellular growth and differentiation are tightly coupled to developmental milestones.

In this study, we investigated the role of mTOR signaling in lens development by generating LE-specific conditional knockouts of *Raptor* (mTORC1) and *Rictor* (mTORC2). *Raptor*/*Rictor* mutants exhibited defective lens morphogenesis, including corneal-lenticular stalk formation and transdifferentiation of the CM into neural retina. These phenotypes correlated with diminished Wnt signaling activity in the lens vesicle and optic cup. Consistent with mTOR-Wnt regulation, pharmacological mTOR inhibition in lens cells downregulated Wnt3 expression, while genetic deletion of *Wnt3* in the LE phenocopied the lens stalk defect, albeit with reduced penetrance. Furthermore, LE-specific ablation of β-catenin or Wnt co-receptors *Lrp5/6* recapitulated the lens herniation observed in *Wls* mutants, implicating autocrine Wnt signaling in maintaining lens integrity. Crucially, all mutants displayed failure of lens vesicle closure—a defect rescued by *Rac1* deletion. Collectively, these results establish an mTOR-Wnt-Rac1 signaling axis that orchestrates cytoskeletal remodeling to drive lens vesicle closure, providing mechanistic insight into the integration of metabolic and mechanical cues during ocular morphogenesis.

## Results

### mTOR signaling is required for lens vesicle separation and ciliary margin specification

To dissect mTOR signaling’s role in lens development, we generated lens ectoderm (LE)-specific double knockouts of *Raptor* (mTORC1) and *Rictor* (mTORC2) using *Le-Cre* (*Le-Cre;Raptor^flox/flox^;Rictor^flox/flox^*; hereafter *Raptor;Rictor^LKO^*), a driver active in the lens placode and persisting in the lens epithelium (Ashery-Padan et al., 2000; Li et al., 2019). At E14.5, *Raptor;Rictor^LKO^* lenses, marked by *Le-Cre*-driven GFP, were severely hypoplastic and exhibited an aberrant fluorescent focus (Fig. 1A, white arrow). Histology identified this structure as a persistent corneal-lenticular stalk (Fig. 1A, black arrow), a hallmark of failed lens-corneal separation during vesicle detachment. Immunostaining confirmed complete loss of mTOR activity in mutants, evidenced by absent phosphorylated mTOR (pmTOR) and its downstream targets pS6 and p4EBP1 (Fig. 1B, dotted lines). Strikingly, *Raptor;Rictor^LKO^* lenses displayed ectopic activation of FGF signaling, with elevated phospho-ERK (pERK) in the anterior lens epithelium—a region devoid of pERK in controls (Fig. 1B, arrows). Moreover, the wild-type pERK gradient (center^high^ in neural retina, periphery^low^ in ciliary margin) was disrupted in mutants (Fig. 1B, yellow dotted lines), correlating with malformed ciliary margins. These findings reveal mTOR’s dual role in suppressing FGF signaling and maintaining retinal patterning, with its loss triggering aberrant signaling rewiring and structural malformations.

The ciliary margin (CM) is organized into three spatially defined compartments: a *proximal zone* co-expressing Sox2 with the neural retina, a *medial zone* marked by Msx1, and a *distal zone* defined by Mitf and Wls (Balasubramanian et al., 2021). In *Raptor;Rictor^LKO^* mutants, Sox2 expression aberrantly expanded into the distal CM, breaching its normal boundary (Fig. 1C, dotted lines), while Msx1 was downregulated. Concurrently, Mitf and Wls became confined to the retinal pigment epithelium (RPE) border, losing their distal CM localization. Structurally, the wild-type CM transitions from a tapered distal edge to a thickened proximal rim, but mutants exhibited uniform retinal-like thickness across the CM (Fig. 1C). This loss of zonation—marked by Sox2 encroachment, Msx1 depletion, and distal marker collapse—collectively indicates transdifferentiation of the CM into neural retina, driven by mTOR-dependent signaling collapse.

**Figure 1.**
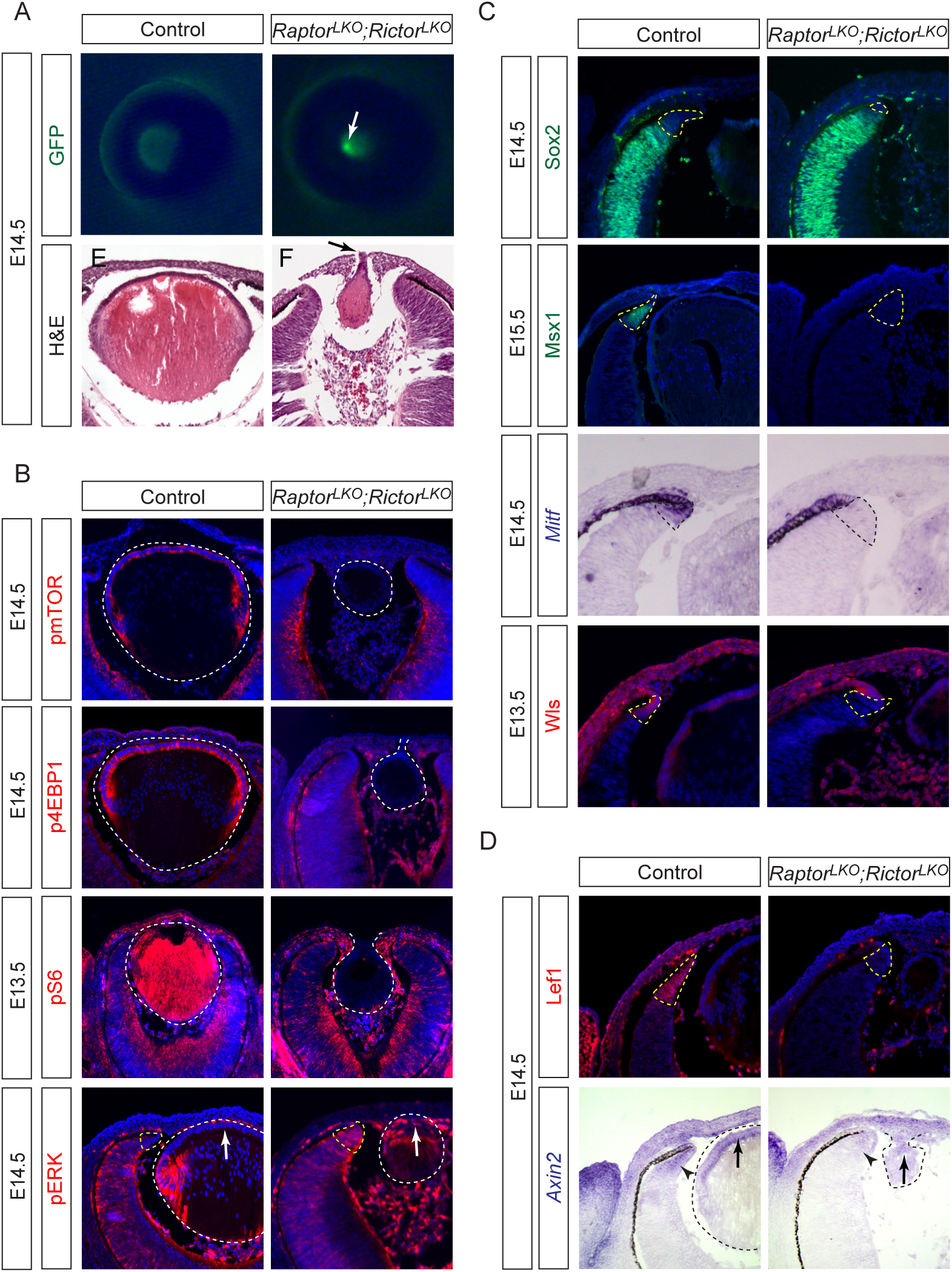
Lens-specific mTOR inactivation disrupts lens-corneal separation and ciliary margin patterning. **(A)** *Raptor;Rictor^LKO^* mutants exhibited hypoplastic lenses with focal GFP expression (white arrow) and persistent corneal-lenticular stalk (black arrow). **(B)** mTOR phosphorylation and downstream targets p4EBP1 and pS6 were absent in *Raptor;Rictor^LKO^* lenses (white dotted lines), but pERK was upregulated in the anterior lens (white arrows) and peripheral retina (yellow dotted lines). **(C)** Ciliary margin markers Msx1, Mitf and Wls were downregulated in *Raptor;Rictor^LKO^* retina, while the neural retina marker Sox2 expanded (dotted lines). **(D)** *Raptor;Rictor^LKO^*mutants showed reduced expression of Wnt signaling response genes Lef1 (yellow dotted lines) and *Axin2* (arrows and arrowheads).

These CM transformations mirror phenotypes observed in optic cup-specific β-catenin knockouts or following genetic inhibition of lens-derived Wnt secretion, strongly implicating Wnt signaling attenuation as a shared mechanism. To test this, we assessed Wnt activity in *Raptor;Rictor^LKO^* mutants via Lef1, a nuclear effector of canonical Wnt signaling. Wild-type embryos exhibit a characteristic rim-to-center Lef1 gradient in the CM, but mutants showed complete loss of this spatial patterning (Fig. 1D, dotted lines). Similarly, expression of *Axin2*—a direct Wnt target and feedback regulator—was drastically reduced in both the peripheral optic cup and lens epithelium (Fig. 1D, arrows). These data demonstrate that LE-specific mTOR ablation non-autonomously disrupts Wnt signaling across the lens-optic cup interface, destabilizing CM zonation. Thus, mTOR functions as a critical upstream modulator of Wnt ligand production or secretion, with its loss uncoupling lens-derived Wnt cues from retinal boundary specification.

### mTOR signaling regulates Wnt expression in the lens

The reduced Wnt signaling observed in both lens and eye cup of *Raptor;Rictor^LKO^* embryos suggests that mTOR signaling acts through both cell-autonomous and non-autonomous mechanisms. This aligns with the potential role of mTOR in regulating Wnt ligand expression, where lens-derived Wnts influence ciliary margin development paracrinally while affecting lens development autocrinally. To investigate mTOR’s direct control over Wnt signaling in the lens, we established primary lens cultures from newborn mice and treated them with Torin, a dual mTORC1/mTORC2 inhibitor. Torin treatment abolished phosphorylation of mTOR and its canonical downstream targets, p4EBP1 and pS6 (Fig. 2A). Consistent with mTORC2’s role in AKT activation, pAKT levels were also suppressed, while pERK remained unchanged. Notably, Torin treatment significantly reduced Wnt3 and Lef1 expression without affecting E-cadherin levels, demonstrating specific regulation of Wnt signaling by mTOR in lens cells.

We next explored the functional role of *Wnt3* in lens development by generating a lens-specific *Wnt3* knockout (*Le-Cre;Wnt3^flox/flox^*, or *Wnt3^LKO^*). At E14.5, 11 out of 52 (21%) of *Wnt3^LKO^* mutants displayed a corneal-lenticular stalk phenotype, marked by a focal GFP signal (Fig. 2B, white arrow), though with lower penetrance than mTOR-deficient models. Histology confirmed persistent adhesion between the lens vesicle and surface ectoderm (Fig. 2B, black arrow), with the stalk expressing surface ectoderm markers E-cadherin and P-cadherin but lacking lens-specific N-cadherin (Fig. 2B, yellow arrow), confirming its ectodermal identity. Notably, *Wnt3^LKO^* lenses showed elevated pERK in the anterior epithelium and stalk (Fig. 2B, arrowhead), mirroring FGF hyperactivity seen in mTOR mutants. While Lef1 expression was reduced in the distal optic cup, residual Wnt activity preserved the ciliary margin’s tapered morphology (Fig. 2B, dotted lines), unlike the complete zonation loss in *Raptor;Rictor^LKO^*. These results demonstrate that mTOR-regulated Wnt3 signaling is necessary but partially redundant for lens ectoderm separation.

**Figure 2.**
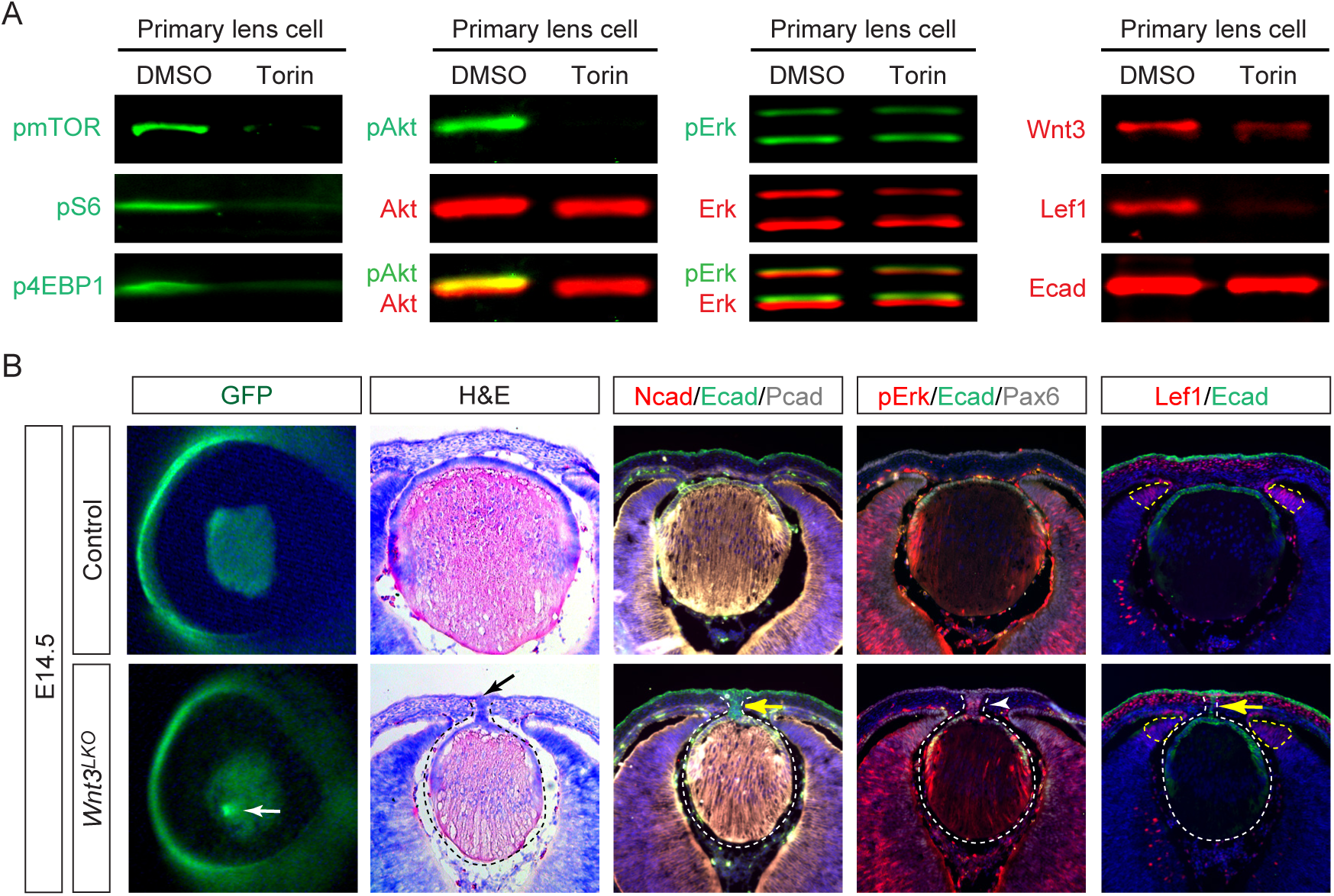
mTOR-regulated Wnt3 signaling is required for lens vesicle closure. **(A)** Treatment with the mTOR inhibitor Torin suppressed pmTOR, p4EBP1, pS6, and pAKT in primary lens culture, while pERK levels remained unchanged. It also significantly reduced Wnt3 and Lef1 levels without affecting E-cadherin. **(B)** *Wnt3* mutant lenses displayed a sharply defined GFP locus (white arrow), confirmed to be a lens stalk (black arrow). The mutant lens lacked N-cadherin but retained ectodermal expression of E-cadherin and P-cadherin (yellow arrow), with upregulated pERK (white arrowhead). Lef1 was downregulated in the peripheral retina.

### Lens-derived Wnt ligands are required for ciliary margin development and lens vesicle closure

The attenuated phenotypes in *Wnt3^LKO^* mutants compared to mTOR-deficient models suggested mTOR regulates additional Wnt ligands beyond Wnt3 in the lens. To test this, we deleted Wntless (Wls), an essential transporter required for secretion of all Wnt proteins (Clevers and Nusse, 2012). Previous studies reported that lens-specific *Wls* deletion resulted in lens hernia, a phenotype more severe than our *Wnt3* knockout, though the underlying mechanisms remained unexplored (Carpenter et al., 2015). At E9.5, *Le-Cre;Wls^flox/flox^*(*Wls^LKO^*) embryos showed normal lens placode induction, with intact Foxe3 expression marking the lens placode (Fig. 3A, dotted lines) and p63 identifying the surrounding non-lens ectoderm (Fig. 3A, arrows). The mutant lens ectoderm also maintained appropriate GFP expression from the Pax6 lens enhancer-driven *Le-Cre* transgene. By E10.5, although reduced Lef1 expression in the peripheral optic cup revealed diminished Wnt signaling (Fig. 3A, arrowheads), *Wls^LKO^* eye displayed normal gross morphology, maintaining robust Foxe3 expression in the lens vesicle, P-Cadherin in the surface ectoderm, pERK in the retina, and Otx2 in the RPE, indicating that early eye development proceeds normally without lens-derived Wnt signals.

By E12.5, *Wls^LKO^* mutants exhibited profound disruptions in ocular development, marked by the transformation of distal RPE to neural retina. This fate shift was evidenced by ectopic N-cadherin expression and loss of P-cadherin at the RPE boundary (Fig. 3B, yellow dotted lines). Concurrently, anterior lens epithelium showed elevated pERK (Fig. 3B, arrows), indicative of hyperactive FGF signaling. Notably, *Wls^LKO^* mutants lacked the usual separation between lens epithelium and surface ectoderm by intervening mesenchymal cells, resulting in direct lens-ectoderm contact (Fig. 3B, solid lines). Laminin staining further revealed breaks in the lens capsule at these contact points (Fig. 3C), culminating in herniation of Maf-positive lens material by E14.5 (Fig. 3D, dotted lines). These findings demonstrate that lens-derived Wnt ligands are essential for both proper lens vesicle closure and ciliary margin development.

**Figure 3.**
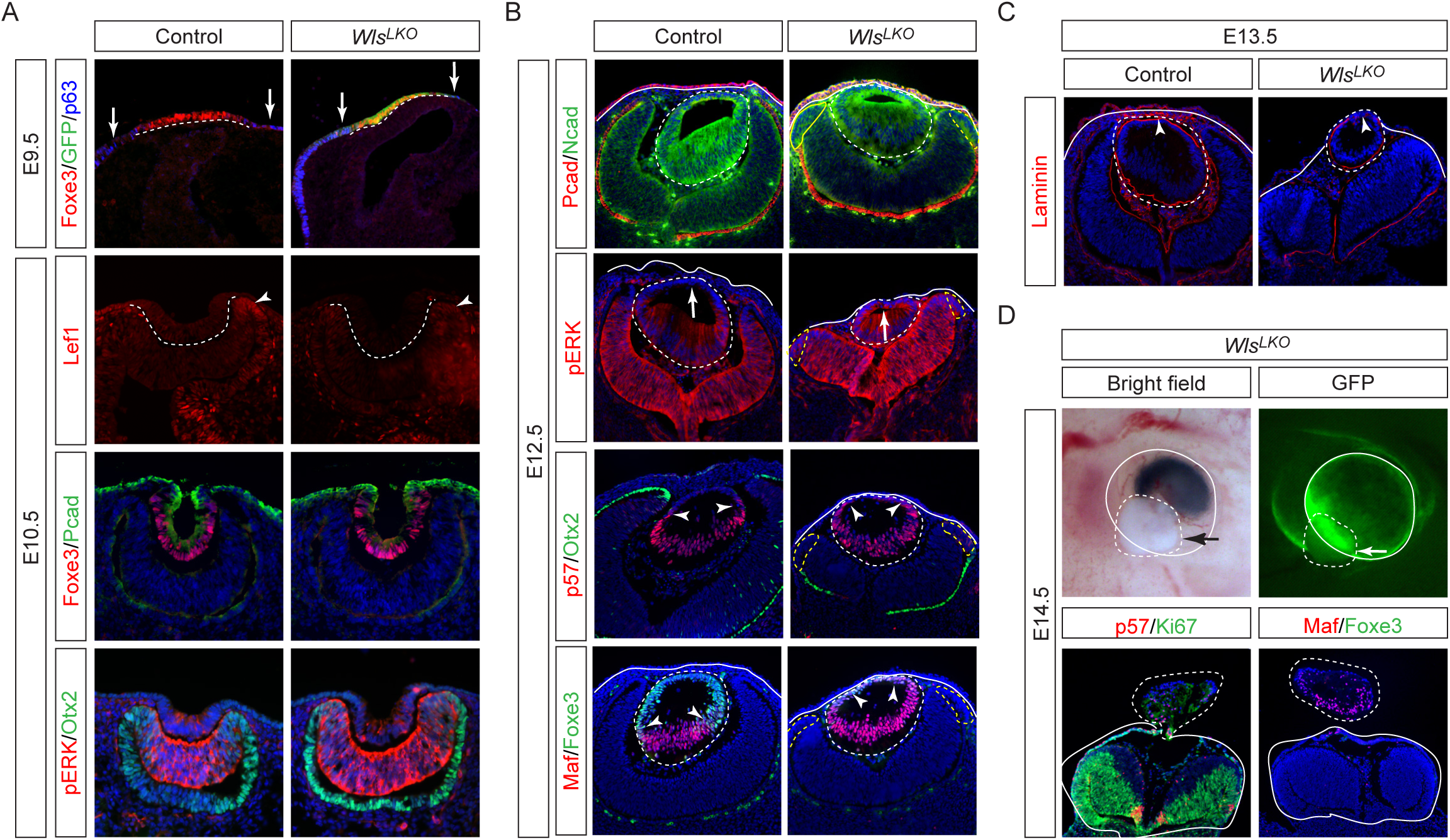
*Wls* deletion causes lens herniation and ciliary margin transdifferentiation. **(A)** E9.5 *Wls^LKO^*mutants exhibited normal Foxe3 and GFP expression in the LE (dotted lines), flanked by p63+ periocular ectoderm (arrows). By E10.5, Lef1 was reduced in the distal optic cup (arrowheads), but the pattern of Foxe3 in the lens vesicle, P-cadherin in the ectoderm, pERK in the retina and Otx2 in the RPE remained unchanged. **(B)** At E12.5, the N-cadherin-expressing lens vesicle abnormally adhered to the P-cadherin-expressing surface ectoderm in *Wls^LKO^*mutants. This was accompanied by pERK upregulation in the anterior lens (arrows), leading to premature induction ofp57 and Maf at the expense of Foxe3 (arrowheads). **(C)** Laminin staining revealed an open lens vesicle in E13.5 *Wls^LKO^* mutants. **(D)** At E14.5, the lens protruded beyond the optic cup in *Wls^LKO^* mutants (arrows), causing expulsion of Maf/p57-expressing lens materials (dotted lines).

### Autocrine Wnt signaling prevents lens hernia

To determine whether lens vesicle closure requires autocrine Wnt signaling, we generated LE-specific double knockouts of Wnt co-receptors *Lrp5/6* (*Le-Cre; Lrp5^flox/flox^; Lrp6^flox/flox^* or *Lrp^LKO^*). At E12.5, *Lrp^LKO^* mutants phenocopied *Wls^LKO^* defects, with direct adhesion between the Foxe3+ lens epithelium and P-cadherin+ surface ectoderm, preventing the normal interposition of mesenchymal cells (Fig. 4A, arrow). Proliferation (Ki67+) persisted in the anterior lens, indicating preserved epithelial homeostasis. By E13.5, the *Lrp^LKO^* lens vesicle expressed lens marker Pax6 and Prox1 in the posterior, but displayed gaps in the Foxe3+/E-cadherin+ anterior epithelium (Fig. 4B). At E14.5, whole-mount imaging revealed lens protrusion through the ocular surface, mirroring *Wls^LKO^* herniation but lacking the severe hypopigmentation indicative of distal eye cup transformation (Fig. 4C). Histology confirmed expulsion of Maf+/Jag1+ lens fibers and a flattened optic cup, phenocopying *Wls^LKO^* structural defects (Fig. 4D). These findings demonstrate that lens-specific deletion of *Lrp5/6* recapitulates the lens hernia defects observed in *Wls^LKO^* mutants without inducing ciliary margin transformation, indicating that the lens hernia phenotype specifically results from disrupted autocrine Wnt signaling within the lens.

### β-catenin mediates canonical Wnt signaling during lens vesicle separation

To assess whether canonical Wnt signaling mediated by β-catenin controls lens structural integrity, we analyzed LE-specific β-catenin knockouts (*Le-Cre;β-catenin^flox/flox^*, or *βcat^LKO^*). In contrast to a contiguous GFP+ corneal ectoderm in E14.5 control embryos, *βcat^LKO^* mutants displayed an expanded GFP+ domain with aberrant fluorescent foci (Fig. 5A, arrowheads), which were confirmed as ectopic lentoid bodies by the lens-specific marker α-crystallin (Fig. 5A, yellow dotted lines). Within the eyes, β-catenin-/α-crystallin+ lenses were hypoplastic and fragmented (Fig. 5A, white dotted lines). These results reveal the dual role of β-catenin in both repressing aberrant lens induction and ensuring proper lens morphogenesis.

**Figure 4.**
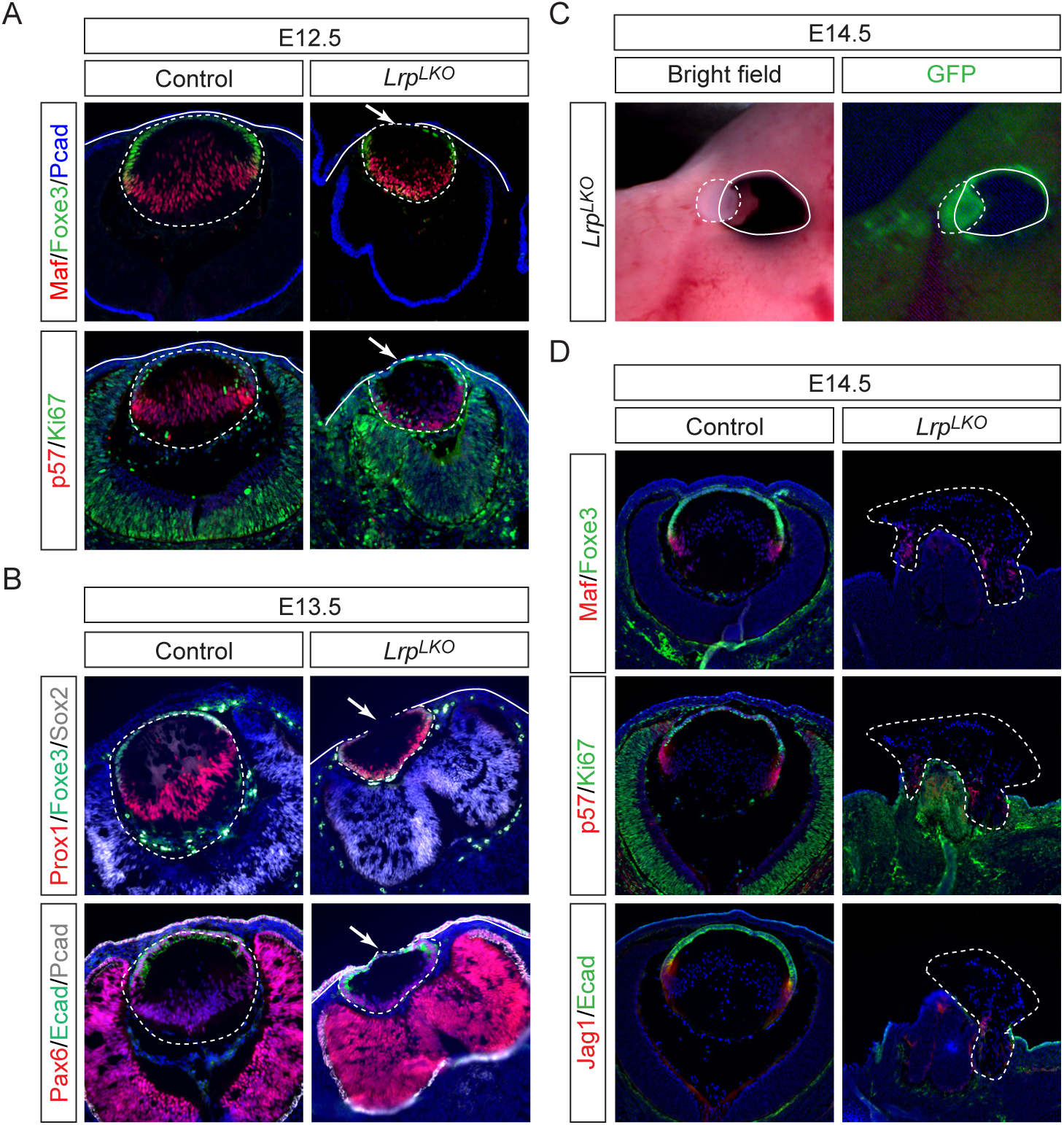
Autocrine Wnt signaling via Lrp5/6 maintains lens integrity. **(A)** E12.5 *Lrp^LKO^*mutants exhibited aberrant attachment (arrows) of the lens vesicle (dotted lines) to the surface ectoderm (solid lines), without disrupting anterior (*Foxe3*+/Ki67+) or posterior (*Maf*+/p57+) lens domains. **(B)** E13.5 *Lrp^LKO^* mutants displayed open lens vesicles marked by a gap in Foxe3/Ecad expression. **(C)** Protrusion of the lens in E14.5 *Lrp^LKO^* mutants. **(D)** In *Lrp^LKO^* mutants, lens materials marked by Maf, p57 and Jag1 expression were expelled by E14.5.

Although lens hernia was not observed in *βcat^LKO^* mutants, we hypothesized that compromised adherens junctions might obscure potential hernia defects. To test this, we utilized a β-catenin allele (*β-catenin^DM^*) that retains adhesive function but lacks transcriptional activity (Valenta et al., 2011). In *Le-Cre; β-catenin^flox/DM^* (*βcat^LDM^*) mutants, we still observed an expansion of the GFP signal from the Le-Cre driver (Fig. 5B, white arrowheads) and occasional intense GFP foci outside the eye (Fig. 5C, yellow arrowheads). At E12.5, the GFP-expressing lens appeared to spill out nasally. Histological sections at E13.5 revealed that the lens, outlined by E-cadherin, maintained a circular shape but remained in direct contact with the P-cadherin-positive surface ectoderm (Fig. 5B, arrows). Further analysis of transverse sections at E14.5 showed that while the mid-eye regions appeared largely normal, the ventral sections revealed lens herniation through the corneal ectoderm (Fig. 5C, dotted lines). Ectopic lentoids were occasionally observed outside the eye (Fig. 5C, yellow dotted lines). Using two antibodies—one recognizing the total β-catenin (βcat^total^) and one targeting the C-terminal epitope missing in β-catenin^DM^ (βcat^CTD^)—we confirmed that adherens junctions remained intact in *βcat^LDM^* mutant lens, as βcat^total^ staining was preserved despite the absence of βcat^CTD^ staining. α-Crystallin staining further confirmed that the irregularly shaped lens had herniated through the corneal opening. These findings demonstrate that canonical Wnt signaling mediated by β-catenin is essential for proper lens vesicle closure and morphogenesis.

**Figure 5.**
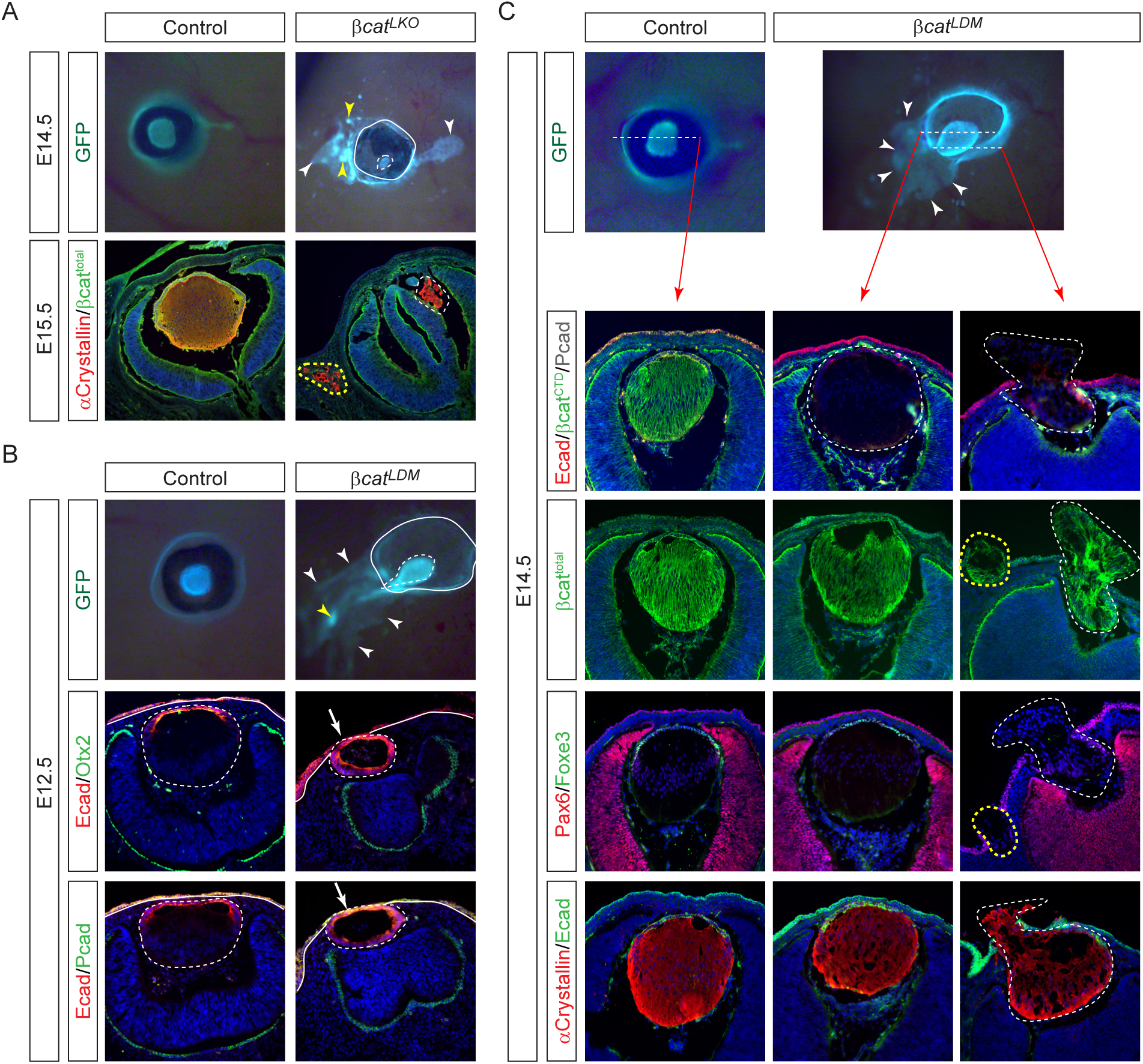
β-catenin plays dual roles in cell adhesion and Wnt-driven lens vesicle closure. **(A)** In *βcat^LKO^*mutants, β-catenin loss resulted in ectopic lentoids (yellow arrowheads) in the periocular ectoderm (white arrowheads). Despite the absence of β-catenin, both ectopic (yellow dotted line) and endogenous (white dotted line) lenses still expressed α-crystallin. **(B)** *βcat^LDM^*mutants, which retain β-catenin’s adhesive function but lack its Wnt signaling role, also exhibited ectopic lentoids (yellow arrowhead) in the expanded GFP+ lens ectoderm (white arrowheads). Although the E-cadherin-expressing lens vesicle remained intact, it remained attached to the P-cadherin-positive surface ectoderm (arrows). **(C)** In *βcat^LDM^*mutants, the lens appeared structurally normal in mid-transverse sections (white dotted lines) but exhibited lens herniation in nasal sections, with ectopic lentoids present (yellow dotted lines).

### Rac1 deletion rescues lens hernia defect caused by Wnt signaling inactivation

To delineate the molecular mechanisms linking Wnt signaling to lens vesicle closure, we performed RNA-sequencing on laser-captured *Wls^LKO^* mutant lenses. Gene Set Enrichment Analysis (GSEA) revealed downregulation of the Wnt signaling pathway, consistent with reduced expression of key Wnt-responsive genes, including *Axin2*, *Dkk1*, *Msx1*, *Fzd7*, *Myc*, *Wls*, *Sfrp2*, and *Tcf7* (Fig. 6B). Interestingly, GSEA analysis also indicated compromised Rho GTPase signaling in *Wls^LKO^* mutants (Fig. 6A), marked by the downregulation of *Rac1gap1*, a highly expressed Rac1-specific GAP, as well as *Arghef2* and *Farp1*, two RhoA-specific GEFs (Fig. 6B), suggesting potential Rac1 hyperactivation at RhoA’s expense. This signature was particularly relevant given our previous discovery that lens morphogenesis requires precise actinomyosin contractility regulated by Rac1-RhoA antagonism (Wu et al., 2024). To test whether Rac1 hyperactivity mediates Wnt signaling’s role in preventing lens hernia, we deleted *Rac1* in *Wls^LKO^* mutants. As previously reported, *Rac1* deletion alone did not alter the overall lens shape at E12.5 but caused a shift from convex to concave lens fiber organization by E14.5, as shown by N-cadherin staining (Fig. 6C). In *Wls^LKO^;Rac1^LKO^*compound mutants, the transformation of the RPE to neural retina persisted, indicated by thickening of the outer eye cup layer and expansion of N-cadherin expression. Strikingly, however, the lens remained within the eye, unlike the lens expulsion observed in *Wls^LKO^*mutants. Thus, *Rac1* inactivation is sufficient to rescue the lens hernia phenotype in *Wls^LKO^* mutants.

Encouraged by the lens hernia rescue in *Wls^LKO^* mutants, we reasoned that if Wnt signaling dysregulation underlies the lens stalk phenotype in mTOR mutants, *Rac1* inactivation should similarly rescue this defect. To test this, we generated *Raptor^LKO^* mutants, which exhibited a lens stalk at E14.5 (Fig. 6D, arrows). Remarkably, the lens stalk phenotype was completely abolished in *Raptor^LKO^;Rac1^LKO^*mutant . Lastly, we deleted Rac1 in the *βcat^LDM^* mutant background. The *Le-Cre; β-catenin^flox/DM^;Rac1^flox/flox^*(*βcat^LDM^*;*Rac1^LKO^*) expanded GFP expression in the periocular mesenchyme with punctuative foci, indicating persistence of ectopic lentoid bodies.

Histological sections confirmed Cre expression in the anterior lens epithelium and loss of βcat^CTD^ staining throughout the lens. However, the mutant lens marked by E- and N-cadherin was completely detached from the P-cadherin-positive surface ectoderm and resided within an intact lens capsule outlined by laminin expression. Taken together, these results showed that mTOR-Wnt-β-catenin axis regulates Rac1 signaling, which is crucial for lens vesicle closure and separation.

**Figure 6.**
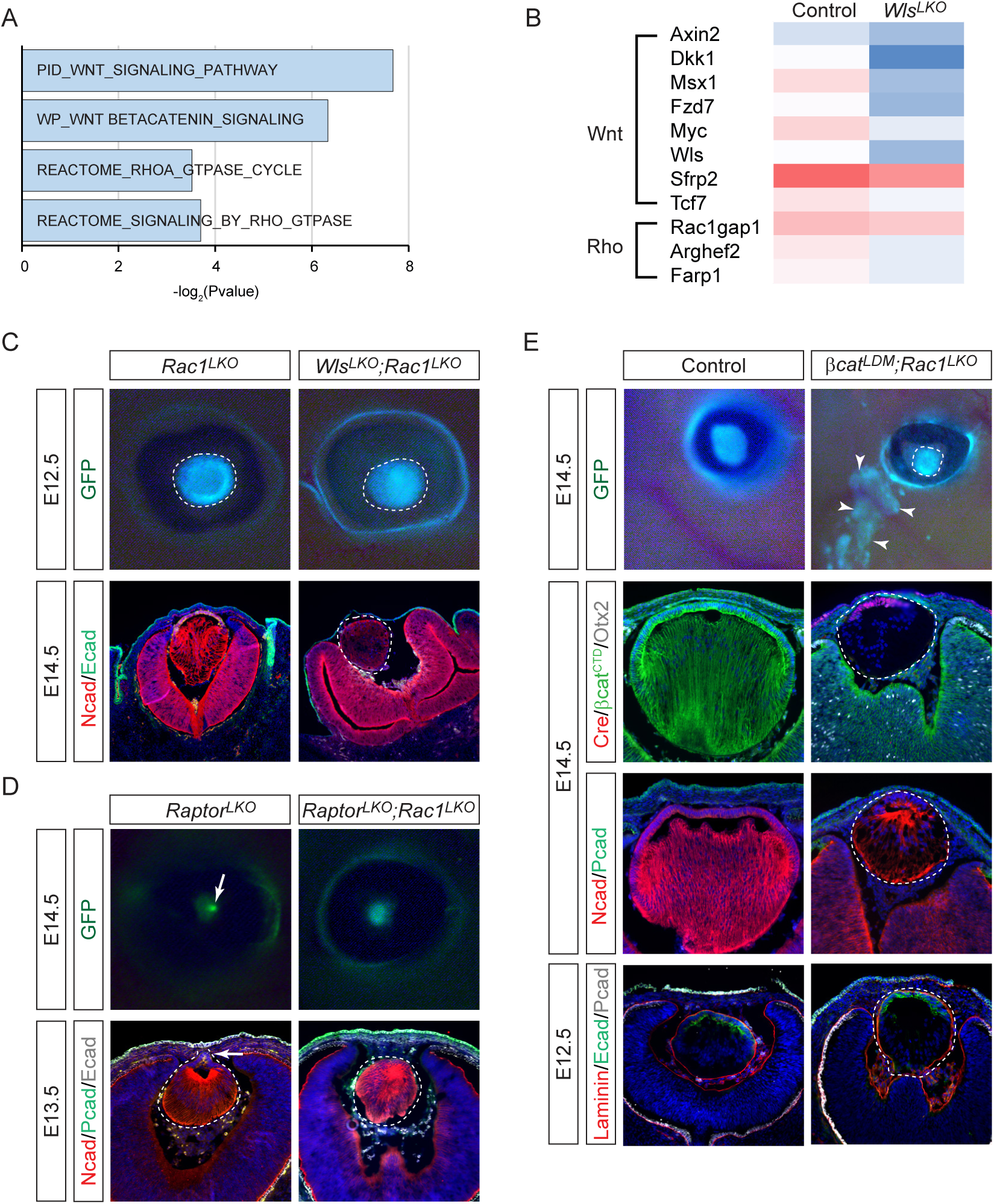
Rac1 inactivation rescues mTOR-Wnt-dependent structural defects. **(A)** GSEA analysis of E11.5 *Wls^LKO^*mutant lens RNA-seq data reveals downregulated Wnt and Rho GTPase pathways. **(B)** A heatmap showed a reduction in Wnt signaling genes and Rho GEFs/GAPs. **(C)** Rac1 deletion altered lens fiber orientation from concave to convex, as shown by N-cadherin expression patterns, while preventing lens herniation in *Wls^LKO^;Rac1^LKO^*compound mutants (dotted lines). **(D)** Rac1 deletion prevented lens stalk formation in *Raptor^LDM^*;*Rac1^LKO^* lens (arrows). **(E)** Despite ectopic lentoid formation outside the eye (arrowheads), *βcat^LDM^*;*Rac1^LKO^* lens (dotted lines) did not develop lens stalks, allowing the interposition of the mesenchymal cells.

## Discussion

In this study, we establish mTOR signaling as a critical upstream regulator of Wnt ligand expression in the lens, coordinating ciliary margin patterning through paracrine signaling and lens vesicle closure via autocrine mechanisms. We identify canonical Wnt/β-catenin signaling as the mechanistic bridge linking mTOR to cytoskeletal dynamics, demonstrating that β-catenin transcriptionally controls the Rho GTPase cycle by modulating expression of key regulatory GEFs and GAPs. Disruption of this pathway induces Rac1 hyperactivation and RhoA suppression, destabilizing actinomyosin contractility required for vesicle closure—a defect rescued by Rac1 inactivation. Strikingly, this Rac1-RhoA imbalance mirrors dysregulation caused by FGF-Abl-p130Cas-Crk signaling in Peters anomaly, revealing convergent erosion of cytoskeletal tension as a unifying pathogenic mechanism (Wu et al., 2024). Our findings redefine Peters anomaly as a disorder of cytoskeletal dysregulation, offering Rac1 as a nodal therapeutic target across genetically heterogeneous cases.

The mechanistic target of rapamycin (mTOR) is an evolutionarily conserved master regulator of cellular metabolism, governing proteomic output by modulating mRNA translation through two key effectors: 4EBP1, which binds and represses TOP motif-containing transcripts, and S6 kinase (S6K), which resolves 5’UTR secondary structures to enhance ribosomal scanning (Laplante and Sabatini, 2012). Exquisitely sensitive to environmental cues such as nutrient availability and oxidative stress, mTOR integrates cellular metabolic status and stress signals to govern cell fate, growth, and survival. Here, we uncover mTOR as a key upstream regulator of Wnt ligand biosynthesis and secretion, revealing a novel layer of crosstalk between nutrient-sensing and developmental signaling pathways. This discovery raises intriguing questions about how maternal nutritional status influences embryonic Wnt-driven morphogenesis, offering potential strategies to mitigate developmental disorders through dietary or pharmacological mTOR modulation. Notably, this mTOR-Wnt axis may have broader relevance: our prior work identified mTOR-mediated control of FGF10 in squamous carcinoma, suggesting a conserved paradigm where mTOR calibrates morphogen signaling across biological contexts (Hertzler-Schaefer et al., 2014). Future studies could investigate mTOR-Wnt crosstalk in adult tissue homeostasis, regeneration, and diverse malignancies, potentially unveiling therapeutic opportunities for diseases rooted in dysregulated growth factor signaling.

Wnt signaling has been extensively studied in eye development. Previous studies have shown that β-catenin deletion results in ectopic lentoid bodies in the periocular ectoderm, while constitutive Wnt activation abolishes lens development, indicating that Wnt signaling suppresses lens fate during early development (Smith et al., 2005). However, interpreting these findings is complicated by β-catenin’s dual role as a component of adherens junctions. To address this, we employed a three-pronged approach to investigate Wnt signaling in early lens development.

First, we ablated Wls, which is essential for Wnt ligand transport from the ER to the plasma membrane for secretion. Second, we deleted the Wnt receptors Lrp5/6 to assess cell-autonomous Wnt signaling. Third, we removed β-catenin to evaluate canonical Wnt signaling. Interestingly, *Wls* mutants did not exhibit ectopic lentoid bodies, suggesting non-lens tissues supply Wnt ligands for early lens fate suppression. In contrast, *Wls* mutants displayed lens hernia and transformation of the ciliary body into neural retina—the latter consistent with previous studies showing that lens-derived Wnt ligands regulate ciliary margin development in a paracrine manner (Balasubramanian et al., 2021). Notably, β-catenin knockouts did not display herniation, likely because adherens junction defects masked Wnt’s closure requirement. By using a β-catenin mutant that lacks Wnt signaling but retains adherens junction function, we successfully recapitulated the lens hernia phenotype, confirming that canonical Wnt signaling is indeed required for lens closure. This highlights the dynamic regulation of Wnt signaling, which must first be repressed and then activated during lens development—a contrast to FGF signaling, which is initially activated for lens induction but subsequently downregulated during lens vesicle closure (Makrides et al., 2022). Such precise temporal coordination underscores how developmental pathways toggle between antagonistic roles to orchestrate tissue morphogenesis.

Peters anomaly is a genetically heterogeneous disorder characterized by corneal opacity, yet its pathogenesis remains poorly understood due to the absence of a unifying mechanistic framework (Bhandari et al., 2011; Nischal, 2012). Clinical presentations diverge between subtypes: Type I involves iridocorneal adhesions, while Type II is defined by lenticulo-corneal adhesions resulting from defective lens vesicle separation. Despite its known developmental origin, the cellular mechanisms underlying Type II Peters anomaly remain elusive. Here, we elucidate a linear signaling cascade in which mTOR-Wnt activation drives aberrant Rac1 activity, identifying Rac1 inactivation as the critical downstream effector required for successful lens vesicle closure. Notably, our recent findings demonstrate that dysregulated FGF-Abl signaling independently causes Peters anomaly, with both genetic and pharmacological Rac1 inactivation rescuing these phenotypes (Wu et al., 2024). The convergence of mTOR-Wnt and FGF-Abl pathways on Rac1-mediated cytoskeletal remodeling reveals previously unrecognized crosstalk during lens morphogenesis. This convergent mechanism not only establishes Rac1 as a central therapeutic target but also redefines Peters anomaly pathogenesis through a shared axis of cytoskeletal dysregulation, offering novel insights into its etiology and treatment strategies.

## Methods and Materials

### Mice

*β-Catenin ^DM^* was obtained from Drs. Jianwen Que (Columbia University) and Konrad Basler (University of Zurich) (Valenta et al., 2011), *Le-Cre* mice from Richard Lang (Children’s Hospital Research Foundation, Cincinnati, OH) (Ashery-Padan et al., 2000), *Rac1^flox^*from Dr. Feng-Chun Yang (Indiana University School of Medicine) (Glogauer et al., 2003), *Rictor^flox^* and *Raptor^flox^* mice from Drs. Markus A. Rüegg and Michael N. Hall (Biozentrum, University of Basel, Basel, Switzerland) (Bentzinger et al., 2008) and *Wnt3^flox^* from Mladen-Roko Rasin (Rutgers University) (Barrow et al., 2003). *β-Catenin ^flox^* (Stock #: 004152), *Lrp5 ^flox^* (Stock#: 026269), *Lrp6 ^flox^* (Stock#: 026267) and *Wls ^flox^* (Strain #012888) were from Jackson lab. Animals were maintained in a mixed genetic background, and at least three animals were analyzed for each genotype. In all conditional knockout experiments, mice were maintained on a mixed genetic background and *Le-Cre* only or *Le-Cre* and heterozygous flox mice were used as controls. All animal experiments were performed according to protocols approved by the Columbia University Institutional Animal Care and Use Committee.

### Histology, Immunohistochemistry, and RNA in situ hybridization

Hematoxylin and Eosin (H&E) staining was performed on paraffin-embedded tissue sections, while immunohistochemistry (IHC) was conducted on both paraffin and cryo-sections following established protocols (Carbe et al., 2012; Carbe and Zhang, 2011). RNA in situ hybridization and immunostaining were performed on the cryosections (10 µm) (Carbe et al., 2013). For immunodetection, the following primary antibodies were used: p57 (#75974) (Abcam); GFP (gfp-1010) (Aves Labs); E-cadherin (#610181), Ki67 (#550609, RRID: AB_393778) (BD Biosciences); P63 (#619001), Pax6 (#PRB-278P) (BioLegend); β-catenin (RRID: AB_11127855, #8480), Cre (#15036), Lef1 (RRID: AB_659971, #2230), N-cadherin (#13116), p4EBP1 (#2855), pERK (#4370), pmTOR (#5536), and pS6 Ribosomal Protein (#5364) (Cell Signaling Technology); Prox1 (PRB-238C) (Covance); Msx1 (RRID: AB_2148804, AF5045), Otx2 (RRID: AB_2157183, AF1979), P-cadherin (RRID: AB_355581, AF761) (R&D Systems); Foxe3 (sc-377465), Jag1 (#6011), Maf (#7866) (Santa Cruz Biotechnology); Wls (RLAB-177) (Seven Hills Bioreagents); Laminin (#L9393) (Sigma-Aldrich); β-catenin (βCTD) (RRID: AB_476831, C2206), Sox2 (RRID: AB_11219471, #14-9811-82) (Thermo Fisher Scientific). Antibodies against α-crystallins were kindly provided by Sam Zigler (National Eye Institute, Bethesda, MD).

For RNA in situ hybridization, probes for *Axin2* (provided by Dr. Guillermo Oliver, St. Jude Children’s Research Hospital, Memphis, TN) and *Mitf* (provided by Dr. Hans Arnheiter, NIH) were used.

### Laser capture microdissection, RNA sequencing and Bioinformatics analysis

Laser capture microdissection and RNA sequencing were performed as previously described (Garg et al., 2020). Preprocessing, quality assessment, and differential gene expression analysis of RNA sequencing data was carried out in R platform. Gene Set Enrichment Analysis (GSEA) was performed using a desktop application developed by the Broad Institute. Collection of annotated gene sets were obtained from the Molecular Signatures Database (MSigDB) v7.2 category C5: ontology gene set. Standard (not pre-ranked) GSEA was performed using normalized counts data imported from *DESeq2* module.

### Cell Culture and Western Blot

Primary lens cells were cultured as previously described (Wang et al., 2017). Briefly, eyeballs were dissected from adult mice (P21 or older), and lenses were carefully extracted with minimal surrounding tissue. Lenses underwent a brief trypsin digestion to remove residual tissue, followed by isolation of the lens epithelium. The epithelium was then digested with dispase and cultured on Matrigel-coated plate in DMEM/F12 medium supplemented with 3% FBS, B27 (Gibco, #17504044), and 5 μM TGF-β inhibitor (Santa Cruz, #SB431542). Lens cells were treated with Torin (150 nM) for 12 hours before western blot analysis.

For immunoblotting, the following antibodies were used: pERK1/2 (Santa Cruz, #sc-7383), ERK1/2 (Cell Signaling, #4695), mouse anti-AKT (#4060), rabbit anti-phospho-AKT (#4060) (all from Cell Signaling Technology), and Wnt3 (Abcam, #ab32249).

## Acknowledgements

The authors thank Drs. Hans Arnheiter, Konrad Basler, Richard Lang, Markus A. Rüegg, Mladen-Roko Rasin, Jianwen Que, Ruth Ashery-Padan, Feng-Chun Yang, Guillermo Oliver, Michael N. Hall, and Sam Zigler for providing mice and reagents. This work was supported by grants from the NIH (R01EY017061 and R01EY025933 to X.Z.). Q.W. is supported by a Pathway to Independence Award (K99EY032171). The Columbia Ophthalmology Core Facility is supported by the NIH Core Grant 5P30EY019007 and unrestricted funds from Research to Prevent Blindness (RPB).

